# The RAG transposon is active through the deuterostome evolution and domesticated in jawed vertebrates

**DOI:** 10.1101/100735

**Authors:** Jose Ricardo Morales Poole, Sheng Feng Huang, Anlong Xu, Justine Bayet, Pierre Pontarotti

## Abstract

RAG1 and RAG2 are essential subunits of the V(D)J recombinase required for the generation of the variability of antibodies and T-cell receptors in jawed vertebrates. It was demonstrated that the amphioxus homologue of RAG1-RAG2 is encoded in an active transposon, belonging to the transposase DDE superfamily. The data provided supports to the possibility that the RAG transposon has been active through the deuterostome evolution and is still active in several lineages. The RAG transposon corresponds to several families present in deuterostomes. RAG1-RAG2 V(D)J recombinase evolved from one of them, partially due to the new ability of the transposon to interact with the cellular reparation machinery. Considering the fact that the RAG transposon survived millions of years in many different lineages, in multiple copies, and that DDE transposases evolved their association with proteins involved in repair mechanisms, we propose that the apparition of V(D)J recombination machinery could be a predictable genetic event.

## Introduction

The recombination-activating gene products known as RAG1 and RAG2 proteins constitute the enzymatic core of the V(D)J recombination machinery of jawed vertebrates. The RAG1-RAG2 complex catalyzes random assembly of variable, diverse and joining gene segments that are present in the jawed vertebrates genomes in numerous copies and together, with hyper-mutation, generate the great diversity of the assembled antibodies and T-cell receptors. Therefore, the RAG1-RAG2 role in the V(D)J rearrangement of antigen receptors is crucial for the jawed vertebrates adaptive immunity (Teng and Schatz 2015). Concerning the origins of RAG1-RAG2, it remains elusive for more than 30 years as the genes were only found in jawed vertebrates (Danchin E. *et al.* 2004). On the other hand, striking similarities between RAG1 and DDE transposase has been noted: common reaction chemistry for DNA cleavage, similar organization of protein domain structure and similarities between recombination signal sequences (RSSs) and terminal inverted repeat (TIRs) targeted by transposases (Kapitonov and Jurka 2005; Fugmann 2010). The hypothetical transposon ancestry of RAG was further supported upon the demonstration of RAG1-RAG2 mediated transposition *in vitro* (Agrawal *et al.* 1998;Hiom *et al.* 1998) and *in vivo* (Chatterji *et al.* 2006; Curry *et al.* 2007; Ramsden *et al.* 2010; Vanura *et al.,* 2007), thought the efficiency of such reactions in vivo is highly disfavored comparing to recombination.

A next step in the understanding of RAG1-RAG2 recombinase evolution was the discovery of a RAG1-RAG2-like locus in purple sea urchin genome, where genes for both proteins are oriented in close proximity in a head-to-head manner as RAG1-RAG2 locus in vertebrates. However this locus lacks TIR and thus does not show the typical features of a transposon (Fugmann *et al.* 2006).Due to the similarity between RAG1 and *Transib* transposon (a family from the DDE transposon superfamily) and the fact that RAG2 lacks similarity to any known transposon protein, even though it harbors Kelch-like repeats and PHD domains as other eukaryotic proteins, led several authors to propose that a *Transib-like* transposon joined the deuterostomian ancestor genome followed by exons shuffling events bringing *Transib* and the ancestor of RAG2 together (Fugmann 2010). As a result, the RAG1-RAG2 locus was then recruited for an unknown function. A second much more recent recruitment as RAG1-RAG1 V(D)J recombinase most likely occurred at the base of the jawed vertebrates evolution. Kapitonov and Koonin (2015) went a step further and provided *in silico* evidence that RAG1 and RAG2 subunits of the V(D)J recombinase evolved from two proteins encoded in a single transposon as they found three sequences that could correspond to fossilized RAG1-RAG2 transposon (including TIRs) in one starfish genome. A major step in the understanding of the RAG1-RAG2 evolution was reported by our group (Huang *et al.* 2016) showing for the first time the presence of an active RAG transposon in the cephalochordate *Branchiostoma belcheri* named ProtoRAG. The full length ProtoRAG transposon is bound by 5 bp target site duplications (TSDs) and a pair of terminal inverted repeats (TIRs) resembling V(D)J recombination signal sequences (RSSs). Between the TIRs reside tail-to-tail oriented, intron-containing and co-transcribed, RAG1-like and RAG2-like genes. The RAG transposon has been recently active in amphioxus as shown by indel polymorphisms. Furthermore the amphioxus RAG1-RAG2-like proteins could mediate TIR-dependent transposon excision, host DNA recombination, transposition and even signal joint formation at low frequency, using reaction mechanisms similar to those used by vertebrates RAGs (Huang *et al*. 2016).

Here we bring more information about the evolution of RAG transposons. We show that beside *B. belchieri,* an active RAG transposon is found in the hemichordate *Ptychodera flava,* that several fossilized transposons are found in several deuterostomes species suggesting that RAG transposon has been active through the history of the deuterostome lineage.

## Results

### Description of an active RAG transposon in *P. flava* and many fossilized transposons in deuterostomes

Due to the discovery of an active RAG transposon in amphioxus *B. belchieri,* we screened all the available genome and EST projects using the query sea urchin RAG1L and RAG2L sequences. Many hits in several deuterostomians species were found, hits are found in protosomians but they show low similarity and correspond to the transib transposons (Panchin and Moroz 2008) and the chapaev transposon family (Kapitonov and Jurka 2007). The family reported by Panchin and Moroz (2008) as well as many other families were found during our survey. However the connection between these families and the RAG1-RAG2 is not clear even if they are related.

Among the hits found in deuterostomes, one of them corresponds to a complete transposon and other several fossilized transposons (see Figure 1 and Supplementary Table 1) in the hemichordate *P. flava.* In other deuterostome species, we found evidence for RAG1L-RAG2L structures without TIRs but with many fragment copies of the RAG1L-RAG2L locus. some species with an incomplete transposon with TIR and RAGL sequences and many other copies of RAG1L-RAG2L fragments. The presence of TIR on many of these copies might indicate that they correspond to fossilized transposons. Transcribed sequences database are available for several deuterostomes and in most of the case RAG1L and RAG2L transcripts are found, complete or incomplete, thus revealing the domestication of the transposon or their activity.

**Figure 1.**
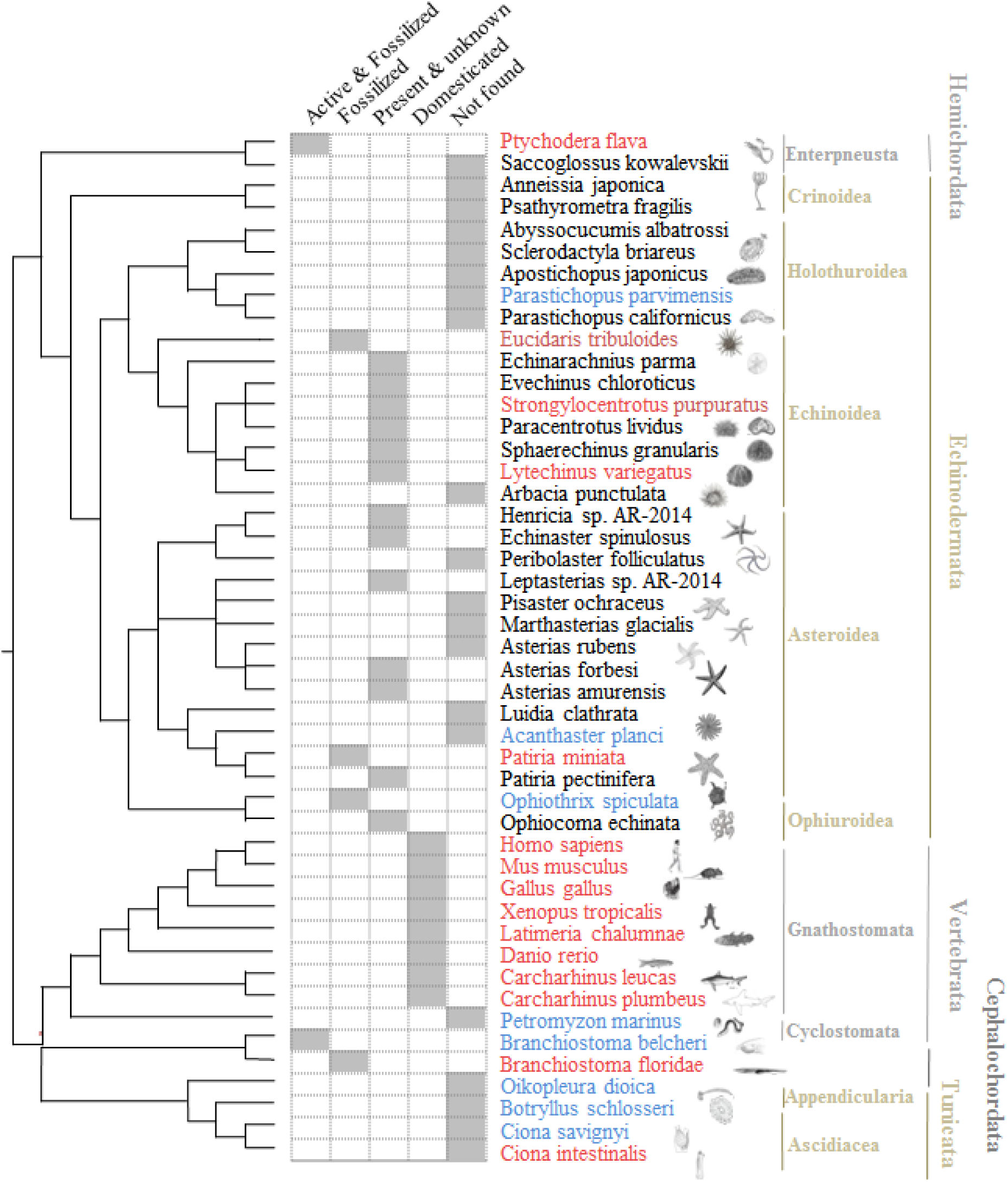
Distribution of the RAG1-RAG2 sequences in deuterostomes. Only species for which the genomic and/or transcription data are available are represented in the phylogenetic tree. Species are colored in red when genomic and transcription data are available, in blue when only genomic data are available and species are colored in black only when expressed sequence data are available.

Based on the phylogeny of the RAG1L and RAG2L protein sequences (see Figure 2 and supplementary table 1 for the phylogenetic analysis and the families description), we can find several RAG families in *P. flava.* Among them, B and C families have unambiguous TIR and TSD structure. Two copies of B family show a TSD-5TIR-RAG1L-RAG2L-3TIR-TSD structure. While one of these two copies encodes a complete RAG1L and RAG2L protein, the other one corresponds to RAG1L and RAG2L pseudogenes. However its presence confirms that the authentic RAG transposon appears in this family. The C family has one copy with TSD-5TIR-(RAG1L-RAG2L)- 3TIR-TSD structure, this copy seems to be inactivated (several in frame stop codons, Supplementary Table 1). In addition, three 5TIR-3TIR copies with no recognizable RAG1/2 genes and one of those copies has both TSD. We also found 12 structures having the 5’ or 3’TIR. We failed to find TSD-TIR structure for other RAG-like families (A and unclassified families) in *P. flava,* this could be due to the poor genome assembly or to the fact that some families have become inactive. Anyway, these findings are sufficient to show that multiple families of RAG transposon have been and are thriving in *P. flava.* Moreover, we found several fossilized transposons in the case of *Patiria minata* as partially described in 2015 (Kapitonov and Koonin 2015), a 5TIR-RAG1L_fragment-3TIR structure containing TSD and no RAG2L protein, a 5TIR adjacent to RAG1L structure (TSD-5TIR-RAG1L) and other several structures having the 5’ and 3’TIR but without internal RAG coding sequences. These structures indicate that RAG was an active transposon during the echinoderms evolution. Afterwards a comparative sequence analysis was made in *B. belcheri, Branchiostoma floridae, P. flava* (Pfl) and *P. minata* (Pmi) TIR sequences (Figure 3) showing no identity between different *Transib*, vertebrate RSS and amphioxus, Pmi and Pfl species except the first CAC nucleotides. Nonetheless both sequences analyzed in amphioxus, share TIR similarity, suggesting a possible common origin of RAG transposon between these two species of amphioxus. However, there is no identity between B and C RAG transposon families in *P. flava,* suggesting, despite the similarity between RAG-like proteins of both families, no TIR similarity between each other, as they may be not reactive or functionally compatible. Previously, an equivalent of RSS nonamer, a stretch of nine highly conserved nucleotides has been found in the amphioxus ProtoRAG TIR, though this ProtoRAG nonamer has no similarity with the nonamer found in RSS (Huang *et al.* 2016). However, there is no such nonamer or equivalently conserved oligomer found in *P. minata* and *P. flava* B and C ProtoRAG family. All this suggests that the nonamer structure is not important in echinoderms and hemichordates phyla, but became important in amphioxus and vertebrates.

**Figures 2.**
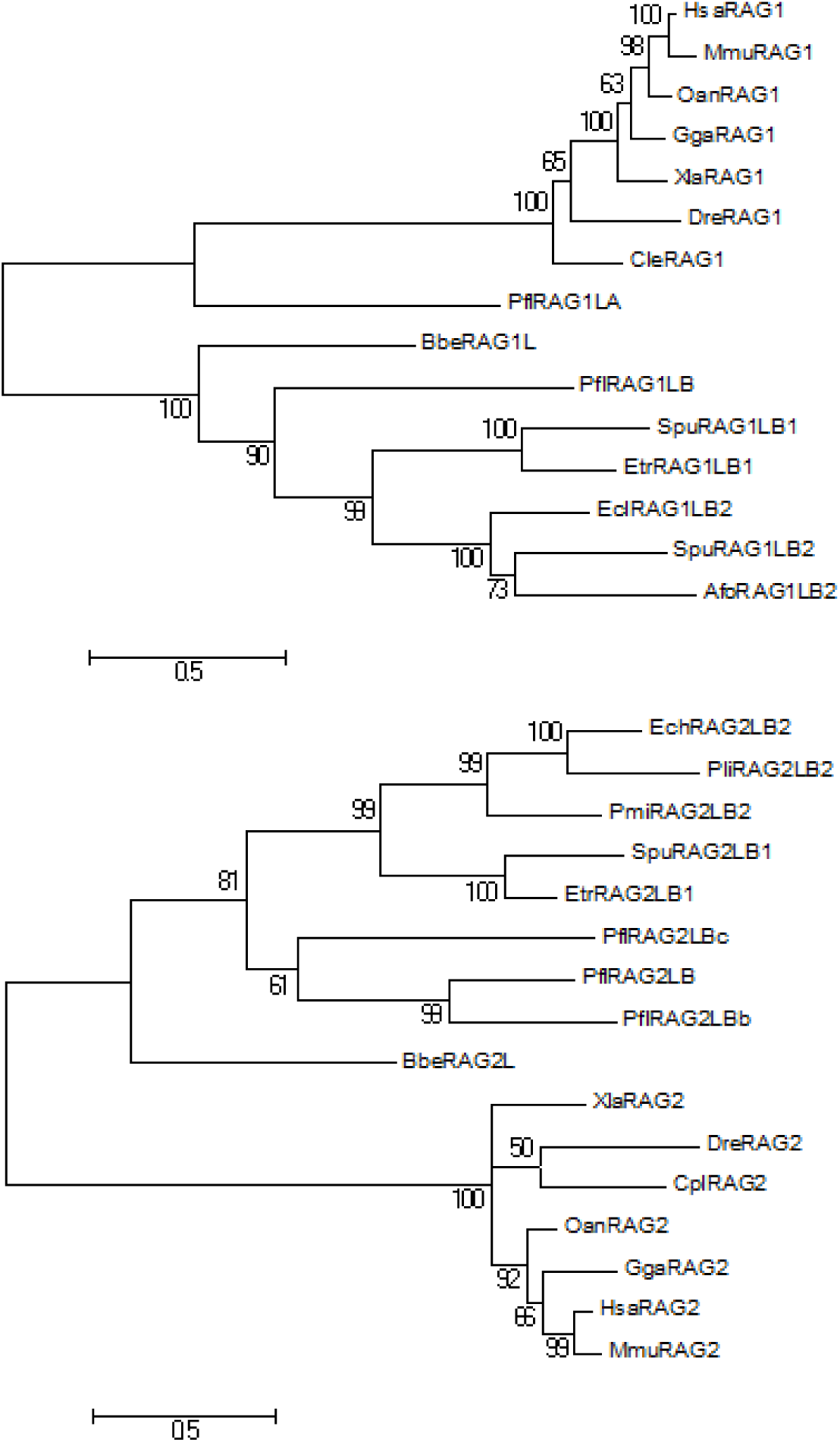
Phylogenetic analysis with the complete RAG1 (2A) and RAG 2 sequences (2B). See also table 1. Phylogeny of two families A and B and other families such as P. flava RAG C are only found in one species (see also table 1). It is therefore difficult to decipher their story.

**Figure 2C.**
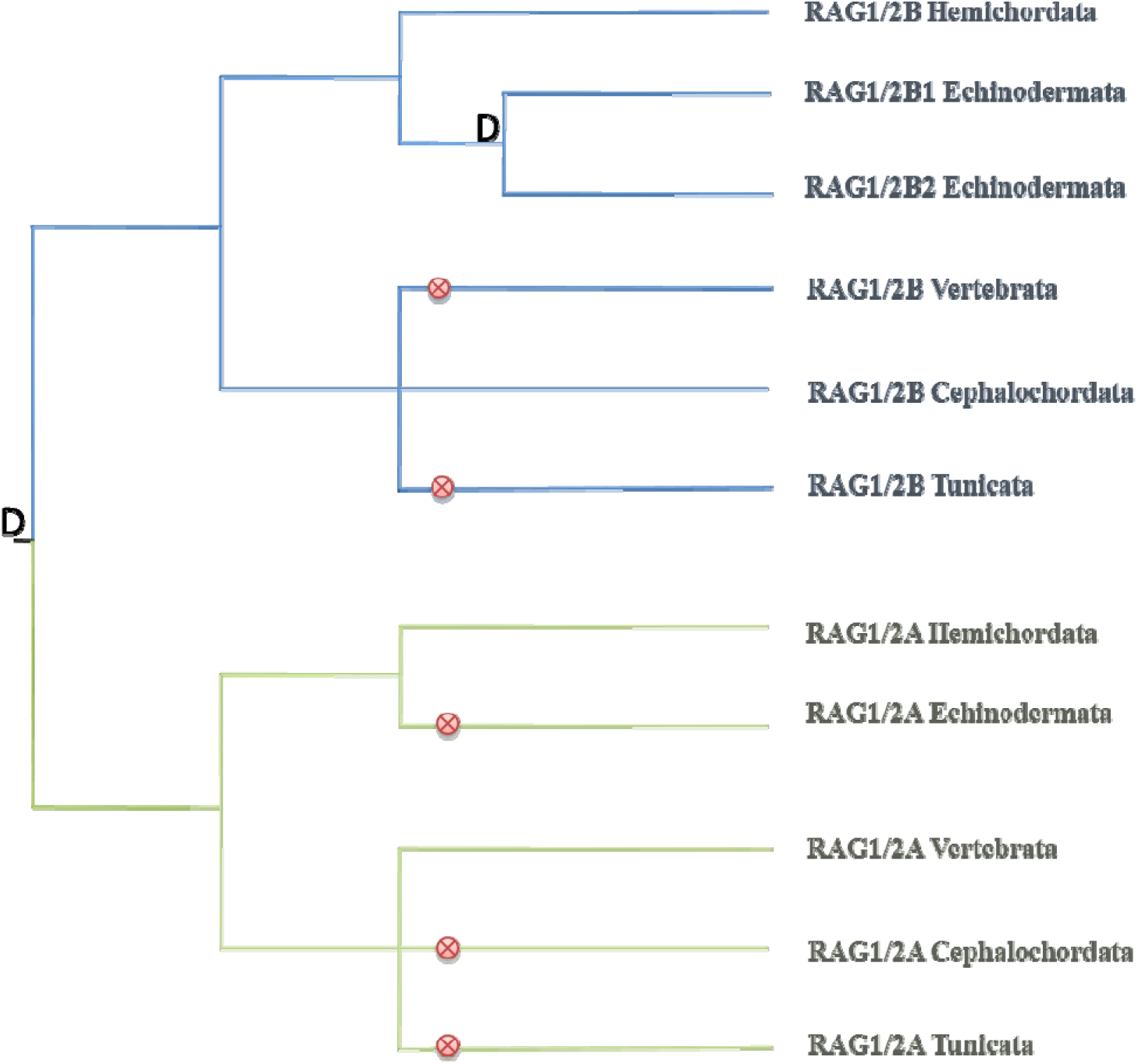
Schematic drawing of duplications and losses of the RAG families during the deuterostome evolution: duplication (D) and lost (⊗).

**Figure 3.**
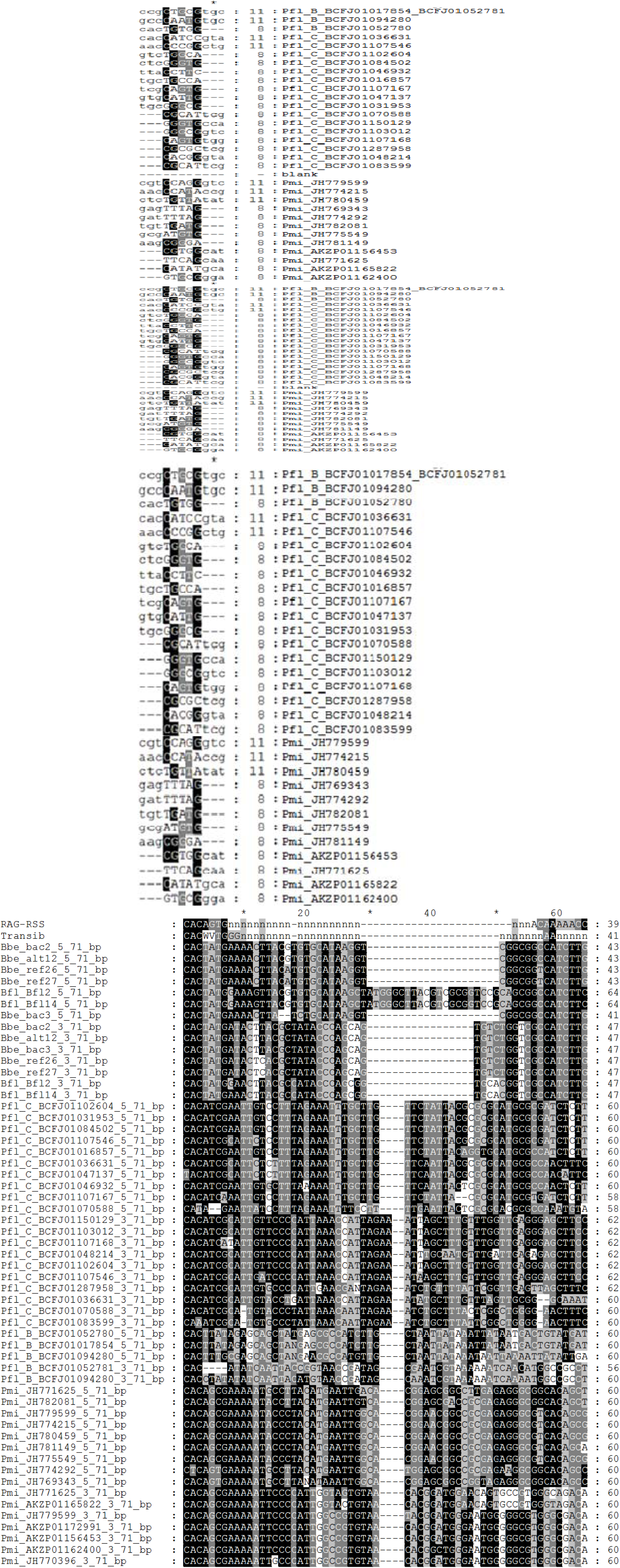
Full length alignment of the TSD and TIR sequences. (3A) Alignment of RAG transposon TSDs and flanking sequences from the *P. flava, P. miniata, B. belcheri, B. floridae* genome. The length of the TSDs is the same: 5bp for the Transib and the RAG-L transposon which indicates similar mechanisms of transposition (3B). Alignment of *ProtoRAG* TIR sequences with the consensus RSS and *Transib* TIR. IUPAC codes used in the alignment: N=A, C, G or T; K=G or T; W=A, T; V=A, C or G Lower case indicates an undetermined nucleotide. Shading indicates sequence conservation, with darker gray indicating a higher degree of conservation. Bbe: *B. belcheri;* Bfl: *B. floridae.* Pfl: *P. flava*, Pmi: *P. minata* RAG *transposon* copy identification numbers correspond to those listed in Table S1.

The species tree in Figure 1 (see also Supplementary Table 1) shows a summary of RAG1L-RAG2L sequences distribution in deuterostomes according to the available data. When genomic and transcription data are available the species names appear in red, whereas when only genomic data are available the species names are shown in blue, and when only available expressed sequence data corresponds to the species name are black. It is likely that the transposon is active if bona fide sequences are present in the genome in several copies and fragments and if the putative transposons are transcribed as in the case for *P. flava* RAGL-B and *B. belcheri.* On the other hand *P. miniata* seems not to be transcribed since only fossilized transposons are found in the genome. In two species of sea urchin, *Eucidaris tribuloides* and *Lytechinus variegatus,* no transcribed sequences are found, but many copies of RAG1L-RAG2L are present on the genome without TIRs indicating that might be fossilized transposons that became inactivated by the loss of the TIR sequences.

The case of *S. purpuratus* is more difficult to understand: the published RAG1L-RAG2L locus (Fugmann *et al.* 2006) renamed here RAG1L-RAG2L B1, was believed to be domesticated, as the RAG1L and RAG2L coding sequences are not interrupted by stop codons, RAG1L and RAG2L are transcribed. And could be functional, but because no TIR sequences has been identified they cannot be a transposon (Fugmann *et al*. 2006). However, we found many fragments which were highly similar to this sequence in the *S. purpuratus* genome, which could reveal a recent transposition event followed by the domestication of one of its copies (see supplementary data and Figure 3). We found another RAGL copy which arose from a duplication event which occurred at the origin of the echinoderms, named RAG1L-B2. The RAG1L-B2 copy is only found fragmented with multiple recent copies in the genome whereas it is complete as RAG1L transcript. A possible explanation for this second locus could be the existence of an active transposon with the genome sequence not well assembled or otherwise a domesticated or recent fossilized transposon. For most of the species we do not have information at the genomic level, but if we find RAGL transcript, this sequence could correspond to an active transposon, domesticated transposon or recent pseudogene. This shows that the transposon has been present in their ancestors.

### Features of the proteins encoded by the RAG-like proteins

In ambulacraria (echinoderm and hemichordate) the deuterostome RAG1-like, 816-1136 aa-long shares around 26.52% sequence identity between RAG1L-B family and vertebrate RAG1, around 33.21% between the orthologous RAGL-A family and the vertebrate RAG1 and only 27.79% between RAG1L-A and RAG1L-B, while inside RAG1L-B family are sharing 48.75% of sequence identity and only 20.13% respect to Transib transposase in terms of core region. As regards to RAG1 lancelet, 30.47% and 37.62% sequence identity are shared with A and B families respectively and only 27.45% with RAG1 vertebrate (see Supplementary Figure 2A). Clusters of high identity are found between RAG1L and vertebrate RAG1 along much of their length, suggesting conservation of multiple functional elements. Vertebrate RAG1 uses four acidic residues to coordinate critical divalent cations at the active site (Ru et al. 2015) and all four are conserved in RAG1L (Supplementary Figure 1A, red highlight). In addition, many cysteine and histidine residues that coordinate zinc ions and play a critical role in proper folding of RAG1 (Kim et al. 2015), are conserved between RAG1L and vertebrate RAG1 (Supplementary Figure 1A, ^*^ and # symbols). However, RAG1L does not share much identity with vertebrate RAG1 in the region corresponding to the nonamer binding domain, consistent with the fact that RAG transposons TIRs have no clear similarity to the RSS nonamer. In fact, different families of RAG1-like have little similarity to each other in the putative nonamer binding domain, consistent with the fact that different ProtoRAG families have very different TIR sequences and no obvious nonamer regions, excluding the amphioxus TIR. Finally, there are also some RAG1-like specific conserved regions (see Supplementary Figure 1, underlined by ^*^). It should be noted that PflRAG1L-A and jawed vertebrate RAG1 show conserved position in the alignment not shared with other RAG1L families, RAG2L 366-535aa long, shares low sequence identity between B family and vertebrate RAG2 (18.69%) and between B family and lancelet RAG2L (25.02%). On the other hand the RAG2L-B family shares around 45.90% while lancelet RAG2L shares only 20.24% identity with RAG2 vertebrate (supplementary Figure 2B). However, the N-terminal six-bladed β-propeller domain (six Kelch-like repeats), which is conserved in both vertebrate RAG2 and ProtoRAG RAG2L, can be discerned in RAG2L. Strikingly, amphioxus RAG2L lacks the entire RAG2 C-terminal region, including the PHD domain as shown previously (Huang et al. 2016). However, this PHD domain is present in all other echinoderm and hemichordate RAG2 proteins (Supplementary Figure 1B). Thus, the absence of this region in amphioxus RAG transposon might be a secondary loss.

### Phylogenetic relation between the RAG families

The phylogenetic analysis with the complete RAG sequences from the available deuterostome species are shown in Figure 2A and 2B and synthesized in Table 1. At least two sub-families have been present in the ancestral deuterostome, named RAGL-B and RAGL-A. Other families such as RAGL-C have not been included in the phylogenetic history as they are found only in one species (Table 1).

**Table 1.**
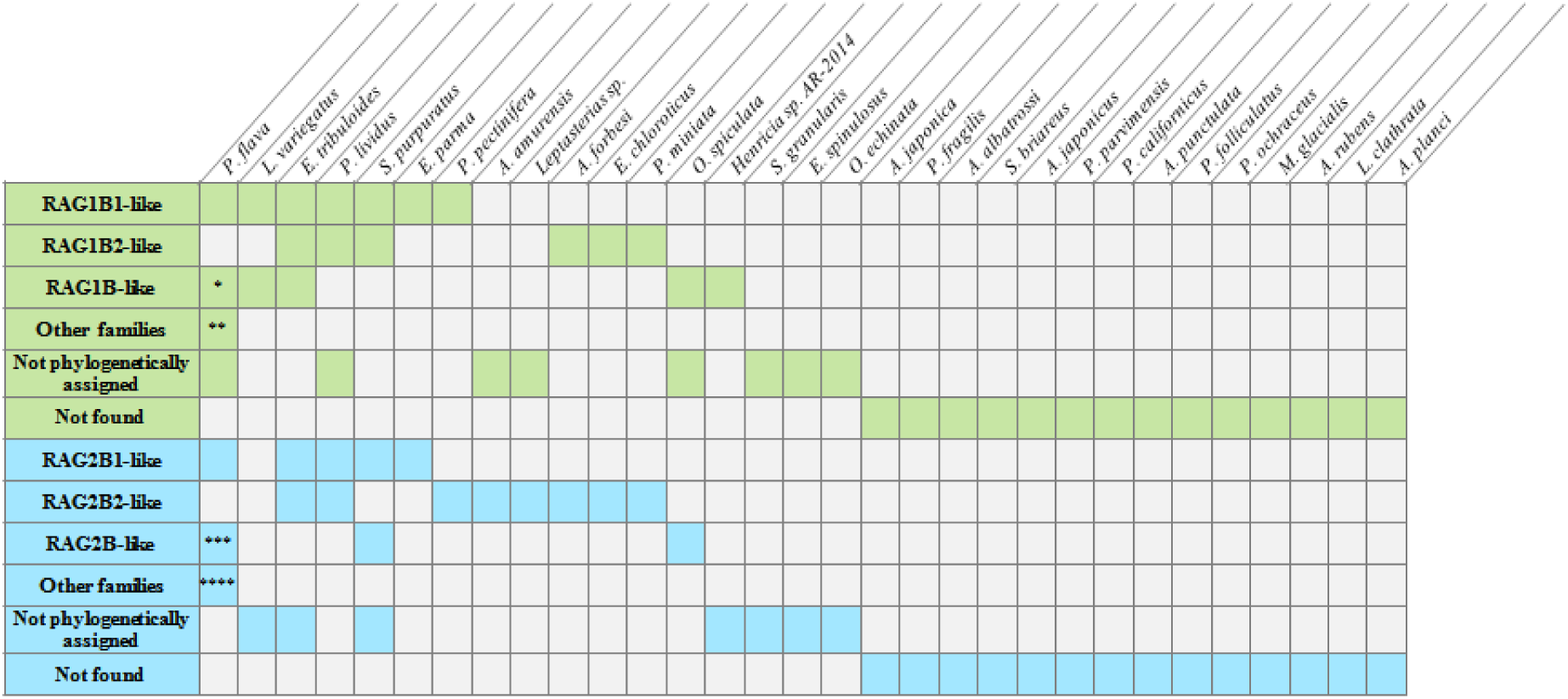
Presence of RAG subfamilies in the different species. Sequences were classified through phylogenetic analysis. Short sequence copies were analyzed one by one against the reference data described in Figure 2A and 2B The classification as B family (or A family labeled with “**”) is straightforward as it is based on orthologous relationships between different phyla (differences between echinoderms and hemichordates for example). Inside B family, two groups named B1 and B2 are found in several echinoderms. If an echinoderm sequence is classified as B family, but not as B1 or B2 we call it B-like (RAG1Bd-like is labeled with “*” while RAG2Bb-like and RAG2Bc-like are labeled with “***”). We have two specific cases, C family only found in *P. flava* (RAG1 labeled with “**” and RAG2 labeled with “****”) and X family in *Ophiotrix spiculata.* The rest of species which do not belong to A or B family, are not phylogenetically assigned due to the fact that none enough phylogenetic signals are available.

In the case of the orthologous relation found between RAG1L-A of *P. flava* (hemichordate) and vertebrates RAG1 recombinase, we can observe that RAGL-A was lost in many lineages excluding hemichordates and jawed vertebrates. RAGL-B conversely, is lost in tunicates and in vertebrates lineage but conserved in several lineages as cephalochordates, hemichordates and echinoderms. Furthermore the phylogenetic analysis shows that RAGL-B has been duplicated in the echinoderms ancestor after the hemichordates/echinoderms split, and both copies have been kept (even if most of them have been inactivated) in most of the echinoderm species (Table 1 and Figure 2C).

### RAG transposon has been active during the deuterostome evolution

From the RAG transposon status: active, fossilized, domesticated, absent (Figure 1 and see the description of an active transposon in *P. flava* and many fossilized transposons chapters), we can proposed the following evolutionary history (Figure 4). The transposon has been active in the deuterostomes ancestor and in the branch that leads to the common ancestor of chordate, still active in cephalochordates and domesticated as a RAG1-RAG2 V(D)J recombinase in the common ancestor of jawed vertebrates. The transposon has been lost in the Petromyzon lineage. The transposon has been active in the branch originated from the node between deuterostomes and ambulacraria antecesors. It remains active in hemichordates inside the subphylum of Enteropneusta (at least on the *P. flava* lineage) but is lost in the other enteropneusts as *S. kowalevskii.* Unfortunately we do not have genome information for the other hemichordates subphyla: Pterobranchia. In the case of the echinoderms lineage, the transposon has been present in the echinoderms common ancestor, in the branch leading to the common ancestor of crinoid, in the clade formed by the sea urchin and holothuroids and in the clade formed by starfishes/ophiures. It has been then lost in the crinoid lineage. The transposon has been active in the branch that goes from the common ancestor of echinoderms to the common ancestor of sea urchin/Holothuroids and starfishes/brittle stars. Concerning the Asteroidea/Ophiuroidea group, the transposon has been active in their common ancestor and has been active in the Ophiure lineage in particular in *O. spicalatus* where the transposon is likely to be active or has lost its activity recently. The transposon seems to have been inactive in the starfish lineage but fragments showing similarities to RAG1-L and/or RAG2-L transposons are found in this species. Furthermore, transposons are clearly found fossilized in *P. miniata*. In the case of sea urchin/holothuroids group, it seems that the transposon has been active in their common ancestor and inactive in the holothurian lineage. We should also note that the transposon seems to be active in some sea urchin lineages as in *E. tribuloide* but much less in others.

**Figure 4.**
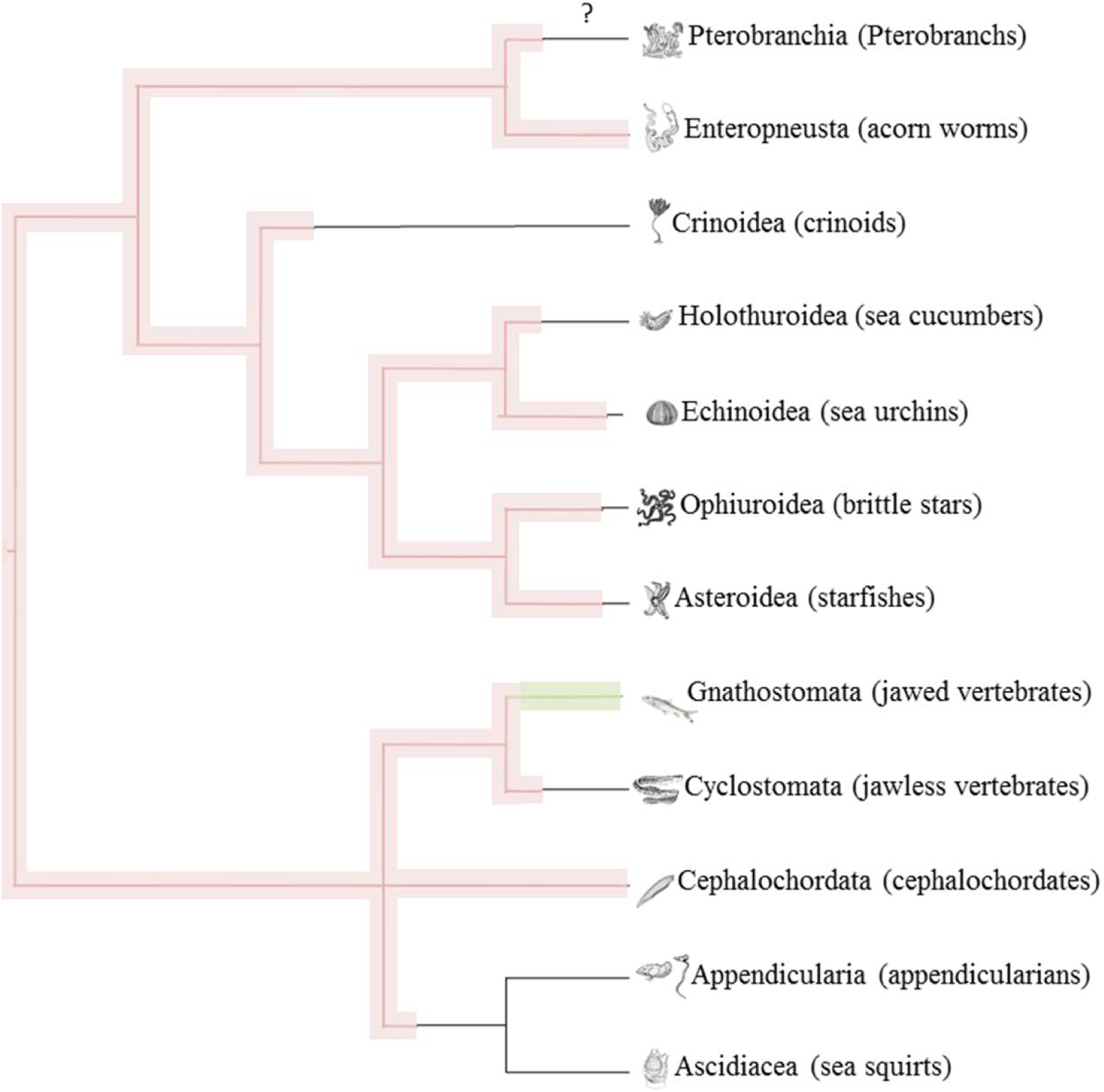
Evolution of the RAG transposon. Transposon activity is indicated in bold pink and V(D)J recombinase activity is indicated in bold green.

## Discussion

In this report we show that a RAG transposon has been present in the deuterostome common ancestors and was active since then in some lineages, fossilized later during evolution and domesticated at least in the case of jawed vertebrates. The structural and regulatory features that cause the jawed vertebrate RAG V(D)J recombinase to favor deletional/inversional recombination over transposition as in the case of the RAG transposase (Huang *et al.* 2016) is not yet resolved. It could be explained by how the cleaved ends and particularly the signal ends are processed. The RAG V(D)J recombinase binds signal ends tightly as excepted for a transposase but it has acquired the possibility to give up these ends efficiently to the non-homologous end joining machinery. This allows recombination and prevents the propagation (Teng and Schatz 2015). Thus the jawed vertebrate V(D)J recombinase differs from the current RAG transposon, as well as its transposon precursor, in how it interfaces with the DNA repair apparatus. This new property occurred likely in the jawed vertebrates common ancestor.

DDE transposases have been shown to interact with repair proteins. For example the Sleeping Beauty transposase interacts directly with the Ku70 repair protein (Izsvák *et al.* 2004) and the pogo transposase of *D. melanogaster* interacts with the proliferating cell nuclear antigen (PCNA), a key protein for DNA replication and repair (Warbrick *et al.* 1998). Therefore, the associations of DDE transposon with DNA repair and replication factors appear to evolve in a convergent manner (Feschotte and Pritham 2007). This characteristic and the fact that the transposon survived during millions of years in multiple copies in different lineages increased the probability of the co-option of the RAG transposon as V(D)J recombinase. Therefore the apparition of V(D)J recombination machinery in the jawed vertebrates phyla could be labeled as a predictable genetic events.

Our results could also explain better the origins of the T-cell receptor and B-cell receptor gene organization. The earlier proposed scenario (Fugmann 2010; Koonin and Krupovic 2015; Hsu and Lewis 2015) involved an insertion of the RAG transposon into the ancestral IG/TCR V-gene, prior to the externalization of the RAG1-RAG2 complex while leaving the RSS-like TIR within the IG/TCR V-gene. This was followed by duplication of this new genetic structure: VRSS-RSSJ. The RAG transposon was then co-opted as V-J recombinase and the system started to work. However, this scenario explains the V-J structure IG light chain, TCR alpha and gamma chain but not the VDJ organization of IG heavy chain or TCR beta and delta chains. Hsu and Lewis (2015) proposed the following scenario for the origin of the D segment: the duplication of the VJ unit, followed by J- to V- recombination and the insertion of non-templated N-region into the signal joint generates a proto D segment. We proposed here an alternative hypothesis to explain the VDJ organization: while one RAG was domesticated (likely RAGL-A orthologue), other RAG transposons (likely RAGL-B orthologue) were still active as one of them split the VRSS-RSSJ copy and gave rise to VRSS-RSSDRSS-RSSJ. RAGL-B transposase became then extinct and finally was lost during vertebrate evolution.

## Material and methods

### Identification of RAG1 and RAG2-like sequence in different data bases

RAG1-RAG2-like locus identified in the echinoderm *Strongylocentrotus purpuratus* and in the vertebrates genome were used as a protein sequence to perform a TBLASTN-based search against the NCBI nr protein, transcriptome shotgun assembly (TSA) and the WGS database as of June 2016 (Altschul *et al.* 1990). These retrieved sequences were extracted and translated by ExPASy Translate tool. Potential open reading frames of RAG1-RAG2 elements used in this study were predicted using FGENESH (Solovyev *et al.* 2006) with the sea urchin organism specific gene-finding parameters. The mRNA sequences were then assembled into contigs by CAP3 (Huang and Madan 1999).

### Phylogenetic analysis

The alignment and trees were constructed using MEGA6 (complete deletion, WAG with Freqs. (+F) correction model, 1000 bootstrap replicates in Tamura *et al.* 2013). Thus, whether they are active, fossilized or domesticated was classified into families. Short sequence copies were analyzed one by one with the reference data set.

### Sequence searches for TIR and TSD motifs

We used three methods to search target site duplication (TSD) and terminal invert repeat (TIR) sequences. In the first method, the upstream and downstream 20 Kb of sequence flanking the RAG1-RAG2-like sequences were extracted and separated into a set of small fragments (using a window size of 60 bp and a step size of 1 bp). In the first method, each upstream fragment was compared with each downstream fragment for 4-6 bp TSDs and possible TIRs using a custom Perl script. We required 40% identity for potential TIR pairs, and allowed only one mismatch for TSD pairs. In the second method, all upstream fragments were compared against all downstream fragments using BLAST. We required a minimum e-value of 100 and sequence identity of 40% in the BLAST search. However, these two methods failed to work well and provided no reliable results. Therefore, we turn to the third method. In this method, we posited that if there are multiple copies of ProtoRAG transposons in the genome assembly, comparison between these copies could help to determine their terminal sequences (TIR, etc.).

We focused on finding more complete elements that contain both TIR and RAG gene fragments, such as “5TIR-RAGs-3TIR”, “5TIR-RAGs” and “RAGs-3TIR”.

Here is our procedure:

1. First we identified all genomic regions containing RAG1/2 fragments by using TBLASTN and the amphioxus and vertebrate RAG1/2 proteins as queries;
2. The region containing RAG1/2 plus upstream 20kb and downstream 20kb was extracted, which we called the RAG region;
3. Because there should have a clear border between the ProtoRAG and the host DNA, we could determine the potential 5’ and 3’-terminal of the ProtoRAG transposon by comparing RAG regions with each other by using BLASTN (see the Figure below);
4. Finally, we examined the potential 5/3-terminal sequences of the RAG regions. Most of them had been destroyed and therefore no detectable TIRs, but several of them shown clear and intact TIR structure.
5. And the TSD if presents, should be right next to the TIR sequences.

Therefore, the sequences containing the RAG1/2-like fragments and the 20 Kb flanking regions were compared to each other and also to the whole genome assembly using BLAST. The terminal sequences were analyzed using a custom Perl script and then subjected to manual inspection.

### Summary of data availability

In order to detect the absence or presence of a given structure in the genome or transcriptome, we need to extract all the available taxonomic information from the NCBI database. It has to be noted that even if the sequence for a given genome is not complete, when RAG1L-RAG2L seems to be an active transposon, we should find an active or at least a fossilized transposons (in several copies). Focusing on the genome database we can find species such as *Parastichopus parvimensis, Acanthaster planci, Ophiothrix spiculata, Petromyzon marinus, Branchiostoma belcheri, Oikopleura dioica, Botryllus schlosseri & Ciona savignyi.* Transcript sequences can be provided for *Saccoglossus kowalevskii, Anneissia japonica, Psathyrometra fragilis, Abyssocucumis albatrossi, Sclerodactyla briareus, Apostichopus japonicus, Parastichopus californicus*, *Echinarachnius parma, Evechinus chloroticus, Paracentrotus lividus, Sphaerechinus granularis, Arbacia punctulata, Henricia sp.* AR-2014, *Echinaster spinulosus, Peribolaster folliculatus*, *Leptasterias sp.* AR-2014, *Pisaster ochraceus, Marthasterias glacialis, Asterias rubens, Asterias forbesi, Asterias amurensis, Luidia clathrata, Patiria pectinifera* & *Ophiocoma echinata.* Finally, together with genomic information and transcript expression we have *Ptychodera flava, Eucidaris tribuloides, Strongylocentrotus purpuratus, Lytechinus variegatus, Patiria miniata, Homo sapiens, Mus musculus, Gallus gallus, Xenopus tropicalis, Latimeria chalumnae, Danio rerio, Carcharhinus leucas, Carcharhinus plumbeus, Branchiostoma floridae* and *Ciona intestinalis*.

## Acknowledgements

We thank the EBM laboratory for advice and Olivier Loison for editing the manuscript.

## Author’s contribution

JRMP, PP and SFH conceived the project and design the study. JRMP, PP and SFH analyzed the results. JRMP, PP, SFH and ALX wrote the manuscript.

The authors declare no competing financial interest.

---

**Figure S1.**
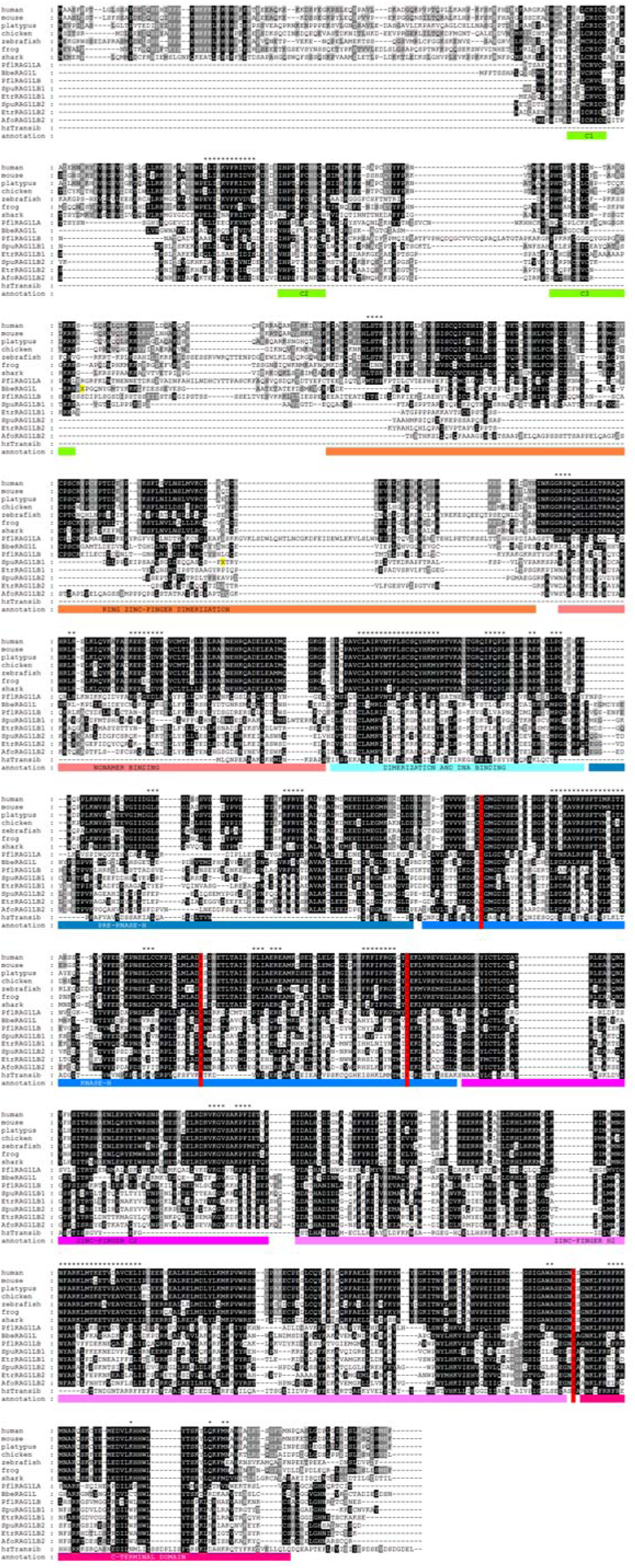
Features of the proteins encoded by the RAG and RAG-like proteins. (A) Protein alignment of RAG1L with vertebrate RAG1. Repeat motifs in amphioxus and the purple sea urchin RAG1L were removed and replaced with an “X” and highlighted in yellow. Three regions of conserved cysteine and histidine residues that might bind zinc are underlined with green bars. The N-terminal zinc binding dimerization domain is underlined with dark-red bars. The subdomains of the RAG1 core region are indicated with colored bars. The conserved acidic catalytic residues are highlighted with red shading (D600, E662, D708 and E962 on mouse RAG1). The PflRAG1LA is more similar to vertebrate RAG1, and those regions were labeled with “*”. GenBank accessions for mouse RAG1, shark RAG1, lancelet RAG2L and sea urchin RAG1L are NP_033045, XP_007886047, KJ748699 and NP_001028179, respectively. (B) Protein alignment of RAG2L with vertebrate RAG2. Color shading shows the conservation of physiochemical properties. The N-terminal amino acid sequences correspond to Kelch-like repeats. The central conserved GG motifs of the six Kelch-like repeats are underlined in red. The plant homeodomain (PHD) is also underlined below the alignment. GenBank accessions for mouse RAG2, shark RAG2, lancelet RAG2L and sea urchin RAG2L are NP_033046, XP_007885835, KJ748699 and NP_001028184, respectively.

**Figure S2.**
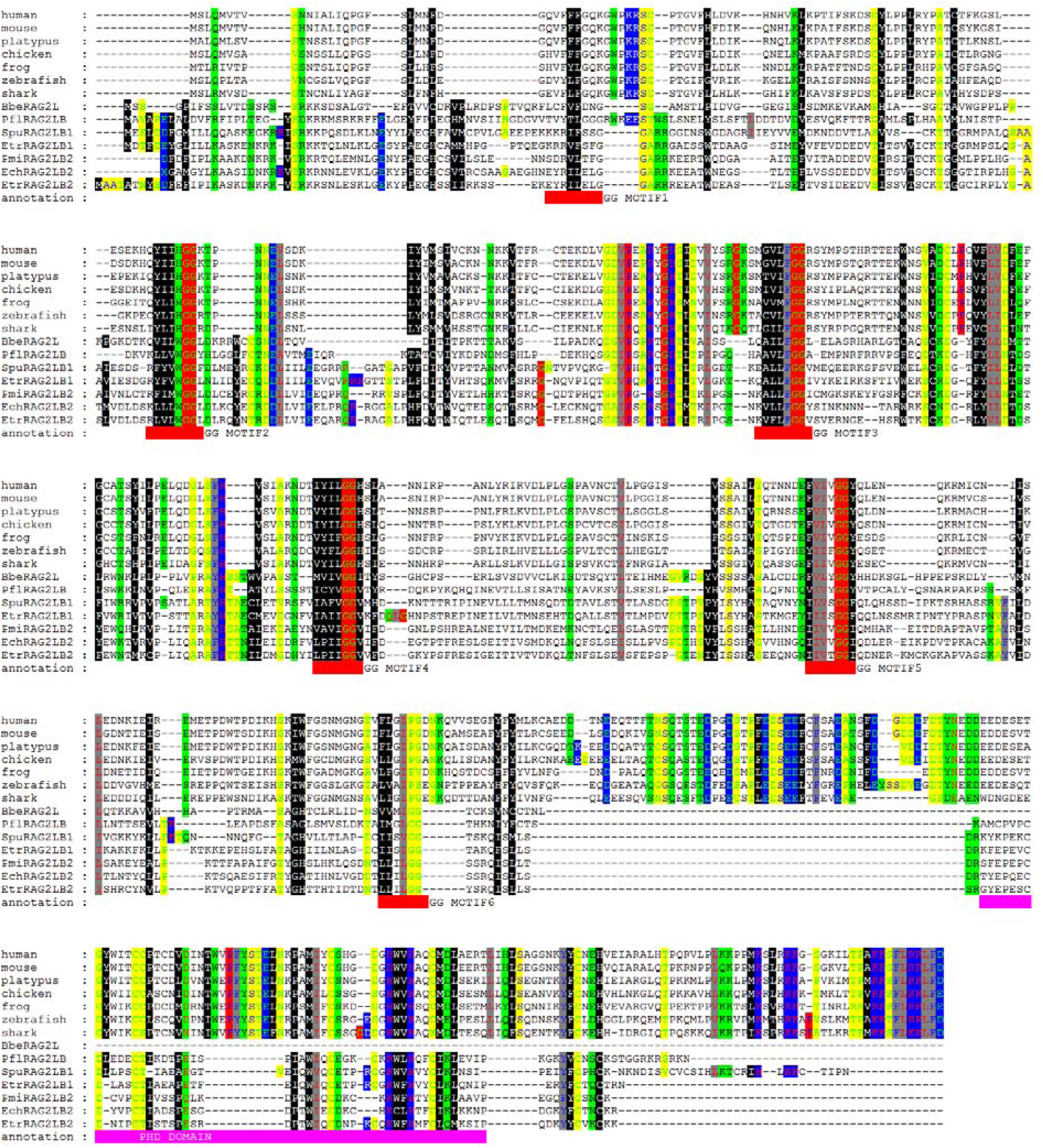
Percent Identity Matrix of RAG1 (S2A) and RAG2 (S2B). In order to provide a multiple alignment, Clustal-Omega requires a guide tree which defines the order in which sequences/profiles are aligned. A guide tree in turn is constructed, based on a distance matrix. Conventionally, this distance matrix is comprised of all the pairwise distances of the sequences. The distance measure Clustal-Omega uses for pairwise distances of unaligned sequences is the k-tuple measure. By default, the distance matrix is used internally to construct the guide tree and is then discarded. By specifying, the internal distance matrix can be written to file.

**Table S1.**
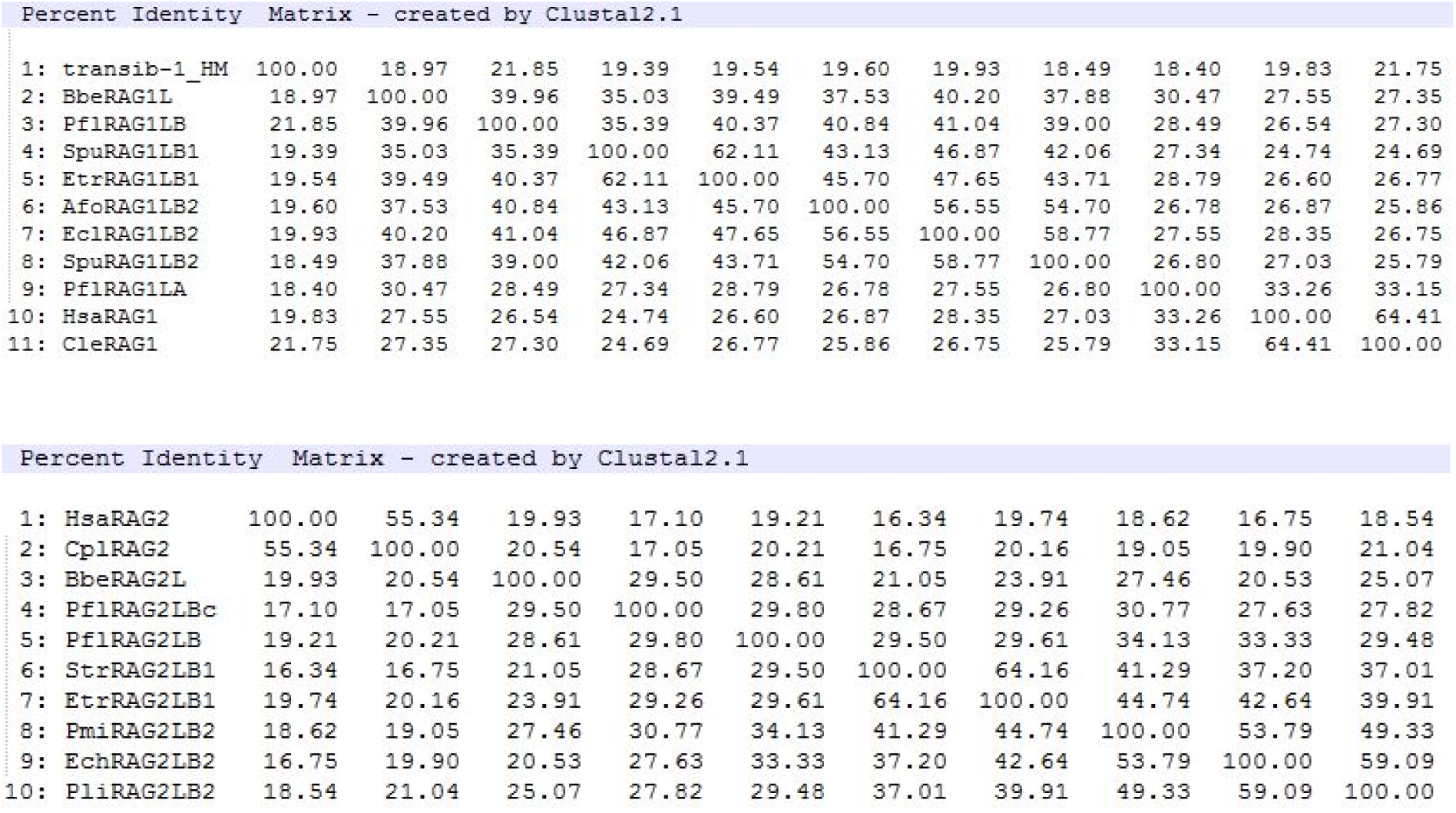
RAGL distribution in non-chordate genome and expressed sequence. Distribution in the cephalordate phyla: *B. belcheri* and *B. floridae* available in ^13^.

| Percent Identity Matrix - created by Clustal2.1 |  |  |  |  |  |  |  |  |  |  |  |
| --- | --- | --- | --- | --- | --- | --- | --- | --- | --- | --- | --- |
| 1: transib-1_HM | 100.00 | 18.97 | 21.85 | 19.39 | 19.54 | 19.60 | 19.93 | 18.49 | 18.40 | 19.83 | 21.75 |
| 2: BbeRAG1L | 18.97 | 100.00 | 39.96 | 35.03 | 39.49 | 37.53 | 40.20 | 37.88 | 30.47 | 27.55 | 27.35 |
| 3: PflRAG1LB | 21.85 | 39.96 | 100.00 | 35.39 | 40.37 | 40.84 | 41.04 | 39.00 | 28.49 | 26.54 | 27.30 |
| 4: SpuRAG1LB1 | 19.39 | 35.03 | 35.39 | 100.00 | 62.11 | 43.13 | 46.87 | 42.06 | 27.34 | 24.74 | 24.69 |
| 5: EtrRAG1LB1 | 19.54 | 39.49 | 40.37 | 62.11 | 100.00 | 45.70 | 47.65 | 43.71 | 28.79 | 26.60 | 26.77 |
| 6: AfoRAG1LB2 | 19.60 | 37.53 | 40.84 | 43.13 | 45.70 | 100.00 | 56.55 | 54.70 | 26.78 | 26.87 | 25.86 |
| 7: EclRAG1LB2 | 19.93 | 40.20 | 41.04 | 46.87 | 47.65 | 56.55 | 100.00 | 58.77 | 27.55 | 28.35 | 26.75 |
| 8: SpuRAG1LB2 | 18.49 | 37.88 | 39.00 | 42.06 | 43.71 | 54.70 | 58.77 | 100.00 | 26.80 | 27.03 | 25.79 |
| 9: PflRAG1LA | 18.40 | 30.47 | 28.49 | 27.34 | 28.79 | 26.78 | 27.55 | 26.80 | 100.00 | 33.26 | 33.15 |
| 10: HsaRAG1 | 19.83 | 27.55 | 26.54 | 24.74 | 26.60 | 26.87 | 28.35 | 27.03 | 33.26 | 100.00 | 64.41 |
| 11: CleRAG1 | 21.75 | 27.35 | 27.30 | 24.69 | 26.77 | 25.86 | 26.75 | 25.79 | 33.15 | 64.41 | 100.00 |

| Percent Identity Matrix - created by Clustal2.1 |  |  |  |  |  |  |  |  |  |  |
| --- | --- | --- | --- | --- | --- | --- | --- | --- | --- | --- |
| 1: HsaRAG2 | 100.00 | 55.34 | 19.93 | 17.10 | 19.21 | 16.34 | 19.74 | 18.62 | 16.75 | 18.54 |
| 2: CplRAG2 | 55.34 | 100.00 | 20.54 | 17.05 | 20.21 | 16.75 | 20.16 | 19.05 | 19.90 | 21.04 |
| 3: BbeRAG2L | 19.93 | 20.54 | 100.00 | 29.50 | 28.61 | 21.05 | 23.91 | 27.46 | 20.53 | 25.07 |
| 4: PflRAG2LBc | 17.10 | 17.05 | 29.50 | 100.00 | 29.80 | 28.67 | 29.26 | 30.77 | 27.63 | 27.82 |
| 5: PflRAG2LB | 19.21 | 20.21 | 28.61 | 29.80 | 100.00 | 29.50 | 29.61 | 34.13 | 33.33 | 29.48 |
| 6: StrRAG2LB1 | 16.34 | 16.75 | 21.05 | 28.67 | 29.50 | 100.00 | 64.16 | 41.29 | 37.20 | 37.01 |
| 7: EtrRAG2LB1 | 19.74 | 20.16 | 23.91 | 29.26 | 29.61 | 64.16 | 100.00 | 44.74 | 42.64 | 39.91 |
| 8: PmiRAG2LB2 | 18.62 | 19.05 | 27.46 | 30.77 | 34.13 | 41.29 | 44.74 | 100.00 | 53.79 | 49.33 |
| 9: EchRAG2LB2 | 16.75 | 19.90 | 20.53 | 27.63 | 33.33 | 37.20 | 42.64 | 53.79 | 100.00 | 59.09 |
| 10: PliRAG2LB2 | 18.54 | 21.04 | 25.07 | 27.82 | 29.48 | 37.01 | 39.91 | 49.33 | 59.09 | 100.00 |

